# Wanting to like: Motivation influences behavioral and neural responses to social feedback

**DOI:** 10.1101/300657

**Authors:** Brent L. Hughes, Josiah K. Leong, Baba Shiv, Jamil Zaki

## Abstract

Human beings revel in social approval and social connection. For example, individuals want to be liked, and frequently surround themselves with people who provide such positive reinforcement. Past work highlights a “common currency” between social rewards like social approval, and non-social rewards like money. But social and motivational contexts can reshape reward experiences considerably. Here, we examine the boundary conditions that deem social approval subjectively valuable. Participants received feedback about their attractiveness from others. Neural activity in reward-related brain structures (e.g., ventral striatum) increased in response to positive feedback, but only when such feedback came from well-liked targets. These heightened reward responses predicted increases in subsequent attraction to well-liked targets. This work suggests that motivational contexts amplify or diminish the value of social approval in a target-specific manner. The value of social approval is thus defined by the extent to which these experiences bring us closer to people we like.

---

Humans are deeply social creatures. We engage in social behaviors that facilitate a number of adaptive social goals, such as forming connections with one another and maintaining a good reputation. What drives our social motivations? Although adaptive goals may drive social behavior, any single social behavior provides only incremental progress toward achieving them. The drive to engage in social acts may instead stem from its proximal, or immediate, subjective value. That is, individuals value cues that signal social connection and social approval because they feel good. The notion that people subjectively value social experiences has spurred a cottage industry on the neurobiology of social reward.

Research in both human and non-human animals provides strong evidence that people value social stimuli much like they value more primary rewards (e.g., food, money). A set of dopamine-rich midbrain structures (e.g., ventral tegmental area) with connections to the striatum (e.g., nucleus accumbens), and areas of cortex (e.g., ventral medial prefrontal cortex; Berridge, 1996; Haber & Knutson, 2010) reliably respond both when we anticipate or experience rewarding outcomes (e.g., tasting food, winning money; Delgado et al., 2000; Tobler, Fiorillo, & Schultz, 2005). Neuroimaging studies have demonstrated that perceiving and engaging with such social content likewise recruits this reward system (Bhanji & Delgado, 2014; Ruff & Fehr, 2014). For example, the reward system responds when people view social stimuli like happy or attractive faces (Aharon et al., 2001; Cloutier, Heatherton, Whalen & Kelley, 2008; Spreckelmeyer et al., 2013), when people receive positive feedback and social approval from other people (Cooper, Dunne, Furey, & O’Doherty, 2013; Izuma, Saito & Sadato, 2008; Korn, Prehn, Park, Walter, & Heekeren, 2012), when people abide by social norms (Stallen, Smidts, & Sanfey, 2013; Zaki, Schirmer, & Mitchell, 2011), and when people engage in prosocial behaviors (Harbaugh, Mayr, & Burghart, 2007; Morelli et al., 2015; Rilling et al., 2002; Tricomi, Rangel,Camerer, & O’Doherty, 2010). Indeed, research that directly compares monetary reward to social reward finds that both classes of stimuli evoke similar neural responses (Ethridge et al., 2017; Izuma et al., 2008; Wake & Izuma, 2017). Our social tendencies may thus be supported by the same reward system that supports our pursuit of other primary needs (Fliessbach et al., 2007; Levy & Glimcher, 2012; Montague & Berns, 2002).

The reward system also predicts behavior towards valued stimuli (Saunders, Richard, Margolis, & Janak, 2017). For example, reward activity predicts subsequent decisions involving other people (King-Casas et al., 2005), changes in real-world behavior (Falk et al., 2015), and population-level outcomes (Genevsky & Knutson, 2015). These findings suggest that the reward system’s responses to social stimuli motivates social decisions and behaviors towards adaptive goals (Preston, 2017).

However, social experiences do not occur in a vacuum (Hughes & Zaki, 2015; Redcay & Warnell, 2017). Just as the value of primary rewards can change (Balleine & Dickinson, 1998; Kringelbach, O’Doherty, Rolls, Andrews, 2003; Hare, Camerer, & Rangel, 2009; Nook & Zaki, 2015; Plassmann, O’Doherty, Shiv, & Rangel, 2008), so can the value of social rewards. Interpersonal contexts and individual motivations can powerfully re-shape the subjective value of social rewards. One reason contextual factors change subjective value is because contexts can change the likelihood that a social cue or behavior will lead to a desired outcome. For example, the specific identity of people around us can differentially activate motives that redefine what is valuable. The value of social approval might increase when it comes from a friend rather than a creep. As such, the value of a social signal (e.g., social approval) may be amplified or diminished in a target-specific manner (e.g., whether the signal comes from a friend or a creep). However, it remains unclear whether interpersonal contexts change the subjective value of social signals and reward-related neural responses.

One context to investigate this question is the experience of social evaluations. People are motivated to maximize their reputation and self-esteem (Baumeister & Leary, 1995; Epley & Whitchurch, 2008; Ryan & Deci, 2000; Taylor & Brown, 1988; Sedikides & Gregg, 2008). As such, individuals seek out and value social approval from others, and frequently surround themselves with people who provide such approval (Aronson & Linder, 1965; Rodman, Powers, Somerville, 2017; Swann, Hixon, Stein-Seroussi, Gilbert, 1990). Indeed, past research shows robust reward signals when people receive positive evaluations about their appearance, personality characteristics, future prospects, and social status (Cooper et al., 2013; Hughes & Beer, 2013; Izuma et al., 2008; Korn et al., 2012; Morelli et al., 2014; Sharot, Korn, & Dolan, 2011; Somerville, Kelly, & Heatherton, 2010; Zink et al., 2008). However, not all positive evaluations may have been created equal. Instead, the specific identify of the source of social evaluations may powerfully re-shape the value of such evaluations. Close and well-liked individuals are more salient and valuable social targets than distant or disliked individuals, and people generally wish to affiliate with well-liked versus distant or disliked individuals (Fareri, Niznikiewicz, Lee, Delgado, 2012; Hughes & Beer, 2012; Hughes, Ambady & Zaki, 2017). Therefore, the value of social approval may be amplified when it comes from well-liked others in comparison to distant or disliked others.

Here, we investigate this question by using a well-validated paradigm used in previous research that elicits the experience of social evaluation (e.g., Gunther Moor, van Leijenhorst, Rombouts, Crone, Van der Molen, 2010; Hughes & Beer, 2013; Rodman, Powers, Somerville, 2017). Participants were told that they were taking part in a study investigating how individuals form impressions of other people. Participants underwent functional magnetic resonance imaging (fMRI) while they rated the likability of peers and received feedback about the extent to which peers liked or disliked them. We examined whether reward-related neural activity in response to interpersonal feedback is amplified or diminished in a target-specific manner (i.e., greater activity after positive versus negative feedback from well-liked as compared to disliked targets). After completing the feedback portion of the task, participants then re-evaluated the likability of each peer. We examined whether reward-related activity in response to feedback would predict subsequent liking differentially for well-liked than disliked targets.

## METHOD

### Participants

22 male participants were recruited in compliance with the human subjects regulations of Stanford University and compensated with $15/h or course credit (mean age = 19.8 years, s.d. = 1.2). Because the cover story described the experiment as being about evaluations of likability and attraction, participants were prescreened to ensure that they were single and heterosexual, as they would be receiving feedback and evaluating their own levels of attraction to pictures of female participants. The sample size was determined a priori to provide power of 0.80 to detect an effect of feedback on self and other ratings, based on an estimated effect size of d = 1.3 derived from recent studies that also employ within-subject, repeated-measures feedback tasks (Cooper et al., 2013; Hughes & Beer, 2013; Korn et al., 2012; Nook & Zaki, 2015; Somerville, 2010). Participants were all right-handed, native English speakers, free from medications and psychological and neurological conditions, and had normal or corrected-to-normal vision.

### Behavioral procedure

#### Task

Participants underwent a well-validated social evaluation task used in previous research (e.g., Gunther Moor et al., 2010; Hughes & Beer, 2013; Rodman, Powers & Somerville, 2017; Somerville, Heatherton & Kelley, 2010). Approximately 1 week prior to the fMRI experiment, participants visited the lab and had their headshot photograph taken. Participants were then led to believe that their photographs would be rated by female peers participating in the study at other study locations, and that participants would have the opportunity to receive feedback from their female peers as well as provide ratings when they returned for the neuroimaging portion of the experiment.

During the fMRI task, participants completed 160 trials of the social evaluation task split evenly across 2 functional runs. Participants first viewed photographs of the female peers ostensibly participating in the task for 3 s and rated the extent to which they were attracted to the target using a 7-point scale, ranging from 1, *unattractive*, to 7, *attractive*. The participant’s rating was highlighted by a green outline on a Likert scale at the bottom of the computer screen for up to 4 s (see Figure 1 for an illustration of the trial structure). On most trials, participants then received feedback from each peer indicating the extent to which the peer found them attractive or unattractive. Feedback from peers was outlined on the Likert scale in red during the final 2 s. Each trial was followed by a jittered inter-trial interval (1—5 s). Although participants believed that the feedback ratings were made by the female target on screen, these ratings were actually randomly-generated. On approximately one quarter of trials, no feedback was displayed. Visual stimuli were presented using Matlab’s Psychtoolbox and projected onto a large-screen flat-panel display monitor that participants viewed in a mirror mounted on the scanner.

**Figure 1.**
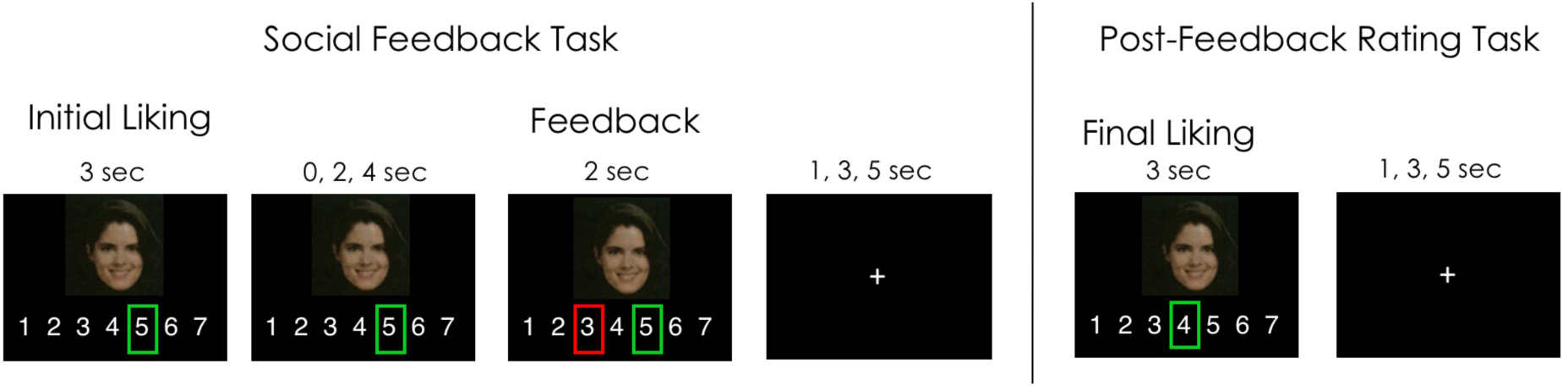
Task Design.

Approximately 5 min after the feedback portion of the task, participants completed 160 trials (evenly split into two functional runs) in which they provided post-feedback ratings of each female peer on the same scale they used on the feedback task. Each rating trial lasted 3 s, and a green outline appeared around participants’ ratings. Unlike the feedback portion of the task, feedback was not presented during these post-rating follow-up trials. Each trial was followed by an inter-trial interval of 1—5 s.

#### Stimuli

The social targets were represented by headshot photographs. Photographs were drawn from the first author’s photo database from prior research and consisted of color pictures of forward-looking faces with neutral expressions. The feedback participants received from the targets depicted in each photograph was randomly distributed across photographs (i.e. such that photograph – feedback pairs were not deterministic).

#### Behavioral analysis

Analysis of behavioral data focused on whether changes in participants’ attractiveness ratings of each face (i.e., post-rating minus initial rating) were influenced by peer feedback. Specifically, we predicted that positive feedback from peers would increase attraction ratings from initial to post-rating, but only when initial attraction to that peer was also positive. To test for the effect of feedback on changes in attraction from initial ratings to post-ratings, we conducted a mixed effects analysis on the trial level using the lme4, nlme and lmerTest packages in R (www.r-project.org; Bates, Maechler, Bolker, & Walker, 2014; Kuznetsova, Brockhoff, & Christensen, 2014; R Core Team, 2014). We computed an impression change score (post-rating minus initial rating) as the response variable. We entered feedback, initial rating, and—crucially—their interaction as fixed effects, and participant as a random effect. Including initial ratings for each trial as a fixed effect covariate ensures that our dependent measures of interest—feedback and its interaction with initial ratings—are not influenced by differences in initial preferences. This approach is also used to control for the potential influence of regression to the mean on analyses of change in ratings. Regression to the mean suggests that social targets that received very high or very low initial attraction ratings should receive re-ratings that are closer to participants’ mean ratings. By including initial ratings as covariates in our mixed effects analyses, we statistically minimize this influence (i.e. the effect of feedback and its interaction with initial rating must explain variance in re-ratings above and beyond differences in initial ratings per se).

### Imaging acquisition and analysis

All images were collected on a 3.0T GE Discovery MR750 scanner at the Center for Cognitive and Neurobiological Imaging at Stanford University. Functional images were acquired with a T2^∗^-weighted gradient echo pulse sequence (TR = 2 s, TE = 24 ms, flip angle = 77°) with each volume consisting of 46 axial slices (2.9-mm-thick slices, in-plane resolution 2.9mm isotropic, no gap, interleaved acquisition). Functional images were collected in 4 runs (each consisting of 160 trials). High-resolution structural scans were acquired with a T1-weighted pulse sequence (TR = 7.2 ms, TW = 2.8 ms, flip angle = 12°) after functional scans, to facilitate their localization and co-registration.

All statistical analyses were conducted using SPM8 (Wellcome Department of Cognitive Neurology). Functional images were reconstructed from k-space using a linear time interpolation algorithm to double the effective sampling rate. Image volumes were corrected for slice-timing skew using temporal sinc interpolation and for movement using rigid-body transformation parameters. Functional data and structural data were co-registered and normalized into a standard anatomical space (2-mm isotropic voxels) based on the echo planar imaging and T1 templates (Montreal Neurological Institute), respectively. Images were smoothed with a 6-mm full-width at half-maximum Gaussian kernel. To remove drifts within sessions, a high-pass filter with a cut-off period of 128 s was applied. Visual inspection of motion correction estimates confirmed that no subject’s head motion exceeded 2.0mm in any dimension from one volume acquisition to the next.

Functional images were analyzed to identify neural activity that was parametrically modulated by initial liking, its interaction with social feedback, and the likability difference score between initial rating and post-rating. Two analytic approaches were used. Both approaches capitalize on estimates of participant ratings and social feedback at a trial-by-trial, within-subject level, by capitalizing on the repeated-measures nature of our experimental design.

#### Whole-brain analysis

The first analytic approach sought to identify whole-brain neural activity associated with parametric increases in initial liking, its interaction with social feedback, and the likability difference scores. All GLMs included six regressors of non-interest modeled participant head movement during the scan. The first GLM consisted of two regressors of interest: the initial liking trial and the social feedback presentation. These regressors were modeled as stick functions at the onset of each trial and convolved with a canonical (double-gamma) hemodynamic response function. In addition, each onset was weighted by 1) the initial rating, 2) the feedback score, and—crucially—3) their interaction. The second GLM included the same regressors of interest, but each onset was instead weighted by the post-rating likability score. We then examined neural activity that parametrically tracked initial liking, its interaction with social feedback, and the post-rating likability scores significantly above baseline (initial rating parametric > baseline; initial rating x feedback interaction > baseline; post-rating likability parametric > baseline). Main effect maps were thresholded at P<0.005, with a spatial extent threshold of k>23, corresponding to a threshold of P<0.05 corrected for multiple comparison (derived with 15,000 Monte Carlo simulations using the current release of the AFNI program 3dClustStim).

#### Volume-of-interest analysis

The second analytic approach involved targeted analyses by specifying independently-defined volumes of interest in Nucleus Accumbens (NAcc) regions associated with valuation and reward in previous research (Knutson & Greer, 2008; Genevsky et al., 2013). Specifically, spherical volumes of interest (8mm diameter) were placed at bilateral foci in the NAcc (MNI coordinates: +/- 10, 14, −6) based on coordinates specified in previous research on valuation and reward (Knutson & Greer, 2008). Activity was spatially averaged within the bilateral NAcc VOIs and then divided by the mean activity over the entire experiment to derive a continuous measure of percent signal changes. Time courses were then shifted 2 volume acquisitions (or 4 s) to account for the hemodynamic lag to peak response per previous research (Knutson et al., 2005). Percent signal change was extracted from trial periods when subjects received social feedback. These were included in the following mixed effects regression models at the trial level, with subjects modeled as random effects: (1) to test the association of trial-by-trial brain activity with initial rating, (2) to test the association of trial-by-trial brain activity with the interaction of initial rating and feedback, and (3) to test the association of trial-to-trial post-rating likability scores with brain activity. Brain activity in any VOI that was greater than 3 standard deviations from the mean percent signal change in that VOI was excluded from regression models. Statistical analyses were performed in R using packages for mixed effects regression analysis (lme4 and nlme).

## RESULTS

### Initial liking and peer feedback influence reward-related neural activity

We first examined parametric trial-by-trial brain activity during the feedback period to identify neural activity that tracked with initial liking ratings as well as the interaction between liking and feedback from peers. Consistent with past work on the reward value of attractive faces (Aharon et al., 2001; Cloutier et al., 2008; Cooper et al., 2013; Spreckelmeyer et al., 2013), we found that activity in striatum (*x*,*y*, *z*: −3, 18, 0, t-stat=3.32, k=53) and other regions (Table I) increased with trial-by-trial liking ratings. Crucially, we found that the interaction between initial liking and peer feedback was also associated with trial-by-trial neural activity in striatum (*x*, *y*, *z*: −9, 12, −3, t-stat=3.68, k=65) and vMPFC (*x*, *y*, *z*: −6, 40, −20, t-stat=3.52, k=49), even after controlling for the influence of initial liking (Figure 2). In particular, activation in these regions tracked trial-by-trial increases in peer feedback, but only to the extent that the participants’ initial liking of peers was also high. To the extent that participants’ initial liking of peers was low, activity in striatum and vMPFC no longer tracked trial-by-trial increases in peer feedback.

**Figure 2.**
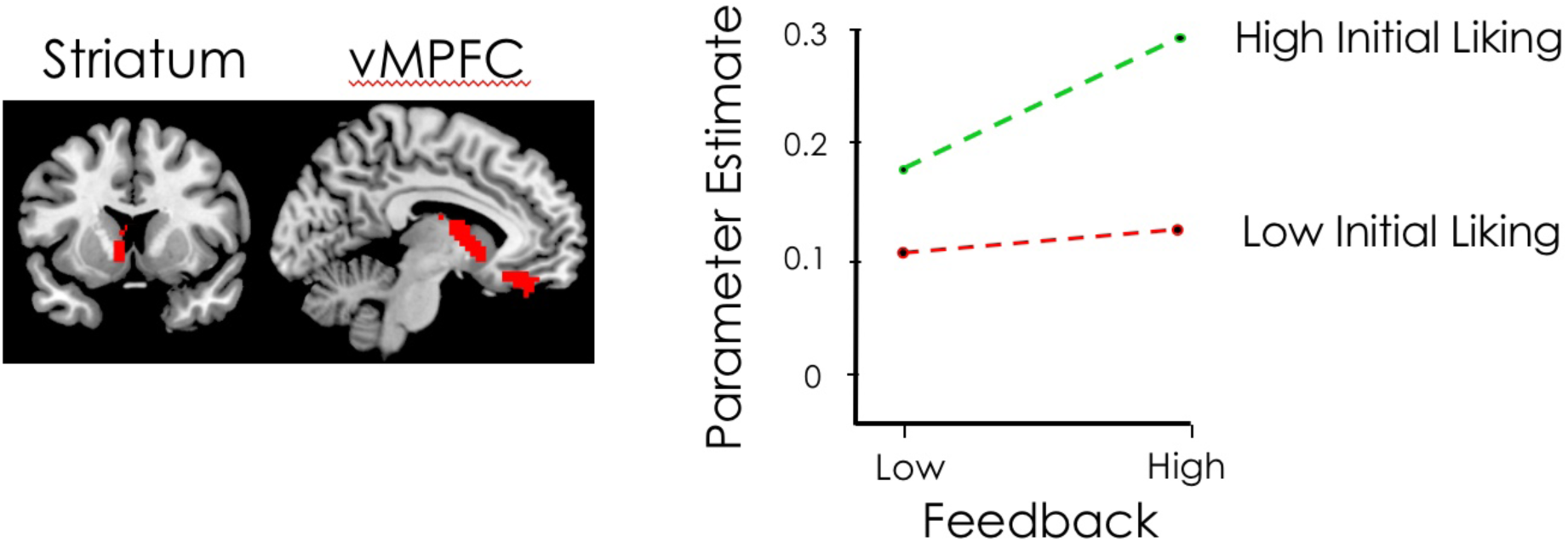
The interaction of liking and being liked on neural responses to social feedback.

**Figure 3.**
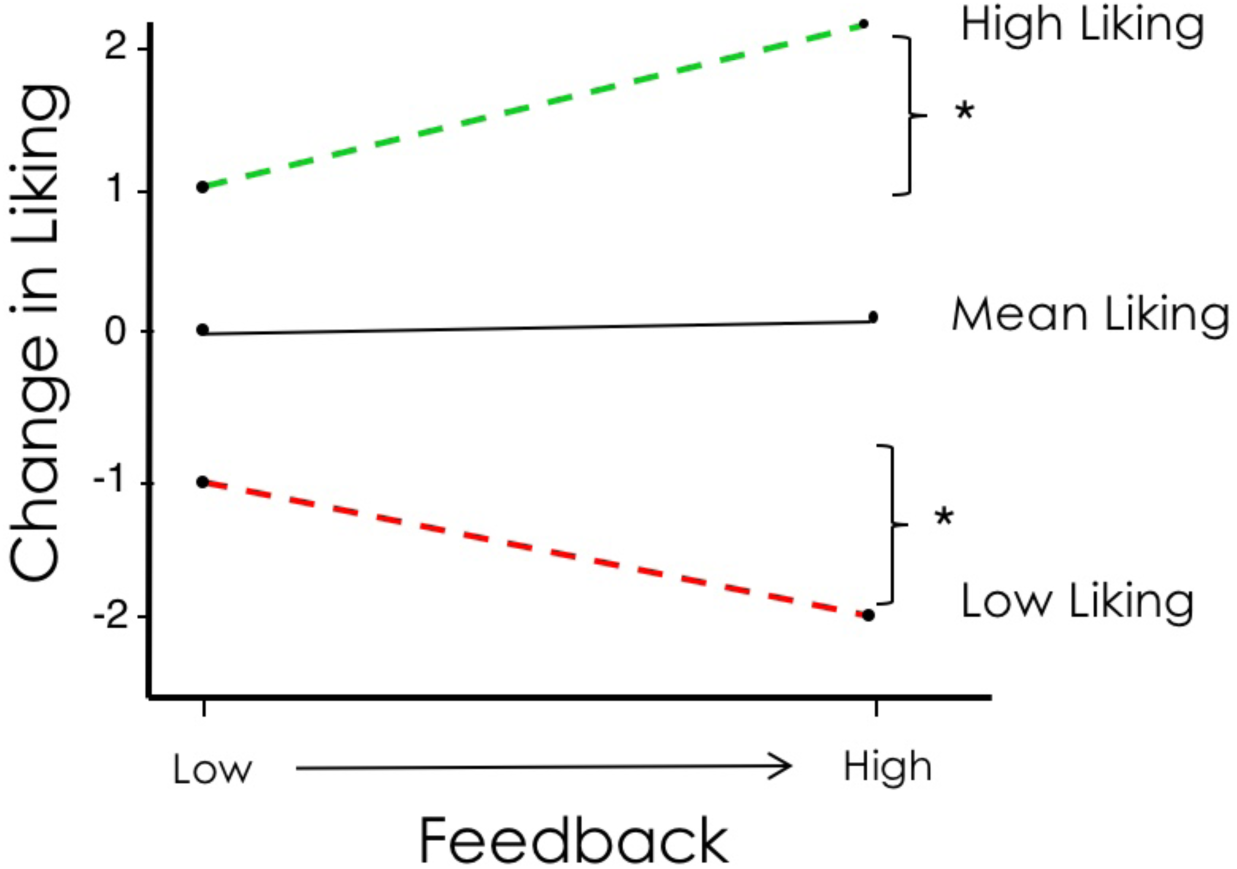
Initial liking and peer feedback predict subsequent liking.

To further test the interaction of initial liking and peer feedback on reward-related neural activity, we examined percent signal changes during the feedback period from independently-defined NAcc ROIs (MNI *x*, *y*, *z*: +/- 10, 14, −6) used in previous research (Knutson & Greer, 2008). These percent signal changes in NAcc were then included in a mixed effects regression model to test the association of trial-by-trial brain activity with the interaction of initial rating and peer feedback. Consistent with the whole-brain effects reported above, we found that initial liking predicted trial-by-trial NAcc activity (b=0.035, t=5.13, p<0.001). Critically, the interaction between initial liking and peer feedback also predicted trial-by-trial NAcc activity (b=0.025, t=2.43, p<0.012). These findings suggest that reward-related activity in striatum tracks the value of positive feedback in a target-dependent manner.

### Initial liking and peer feedback predict subsequent liking

We then examined whether participants changed their impressions of a peer after receiving feedback from them. We computed a likability difference score between post-ratings from initial ratings to quantify whether participants enhanced impressions of a peer (i.e. a positive difference score) or reduced impressions of a peer (i.e. a negative difference score) following feedback from them. To test these effects, we computed a trial-level mixed effects regression model predicting likability difference scores from initial likability ratings, peer feedback, and the interaction between initial ratings and feedback, with subject as a random effect. This model revealed a significant interaction between initial liking and peer feedback (b=0.03, t=3.31, p<0.001), controlling for initial liking ratings (Figure 4). Participants enhanced impressions of a peer following positive feedback, but only when initial liking for that peer was also high. When initial liking for a peer was low, participants reduced impressions of a peer following positive feedback from them. Hence, positive feedback led to enhanced impressions of peers, but only when participants initially liked a given peer. This suggests that the consequences of positive feedback on liking and the desire for social connection might critically depend on the source of the feedback (e.g., a liked or disliked peer).

**Figure 4.**
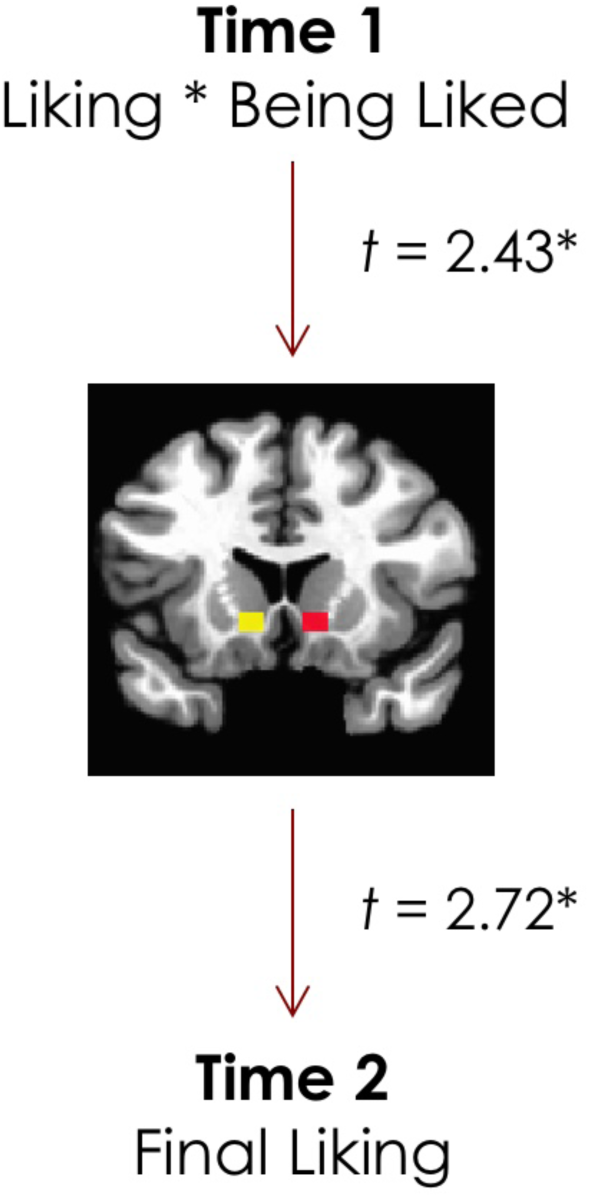
Pathway from being liked to liking.

### Reward-related neural activity predicts subsequent liking

Finally, we tested whether reward-related neural activity at the time of receiving feedback from peers predicts subsequent liking. To do so, we examined parametric trial-by-trial whole-brain activity during the feedback period to identify neural activity that predicted the post-rating likability scores. We found that trial-by-trial neural activity in striatum (*x*, *y*, *z*: −12, 20, - 10, t-stat=5.20, k=102) and vMPFC (*x*, *y*, *z*: 3, 48, −18, t-stat=3.42, k=18) also predicted increases in subsequent liking. To further test this effect, we examined percent signal changes during the feedback period from independently-defined NAcc ROIs (MNI *x*, *y*, *z*: +/- 10, 14, −6; Figure 4) used in previous research (Knutson & Greer, 2008). These percent signal changes in NAcc were then included in a mixed effects regression model to test the association of trial-by-trial brain activity with subsequent liking ratings. Consistent with the whole-brain effects, we found that trial-by-trial NAcc activity predicted increases in subsequent liking of peers (b=0.16, t=2.72, p<0.05). These findings suggest a mechanism by which being liked leads to liking: reward-related activity in ventral striatum tracks the target-dependent value of positive feedback, which predicts changes in subsequent liking (Figure 4).

## DISCUSSION

Here, we show that social and motivational contexts can reshape the subjective value of social rewards, such as signals of social approval. We used a well-validated social evaluation task to probe whether reward-related activity responds to interpersonal feedback in a target-dependent manner. When participants initially liked a given target, they later reported *more* attraction when these well-liked targets provided positive feedback. When participants initially disliked a given target, they later reported *less* attraction when these disliked targets provided them with positive feedback. These effects were mirrored in reward-related brain structures: activity in ventral striatum—defined both at whole-brain and neuroanatomical levels—increased in response to positive feedback, but *only* when such feedback came from well-liked targets. These heightened reward-related responses then predicted increases in subsequent attraction to well-liked social targets. Our findings reveal a framework by which social evaluative processing shapes social preferences: the value people place on positive feedback from others critically depends upon the source of that feedback, which in turn shapes the value associated with social targets.

These findings contribute to a growing body of research on social evaluative processing, and suggest a mechanism by which people enhance themselves and others. People are motivated to feel good about the self and to socially connect with other people (Dunning, Heath, Suls, 2004; Kunda, 1990; Sedikides & Gregg, 2008; Taylor & Brown, 1988). As such, people value and seek out social approval: it accomplishes reputational goals as well as signals progress towards social connection (Leary, Tambor, Terdal, Downs, 1995; Swann, Pelham, Krull, 1989). Indeed, past work shows that people value social approval from peers, and exhibit greater reward-related activity in response to such signals (Cooper et al., 2013; Gunther Moor et al., 2010; Guyer et al., 2008; Izuma et al., 2008; Korn et al., 2012; Morelli, Torre, & Eisenberger, 2014; Somerville, Kelley, & Heatherton, 2010). Our findings conditionalize these insights, by showing that reward activity increases in response to positive feedback, but only when such feedback comes from well-liked targets. These heightened reward responses then predict further increases in subsequent attraction to well-liked social targets. This suggests that the self-enhancement goals may interact with social connection goals, and in doing so shape both the value of interpersonal feedback and the value of social targets. People may not value positive feedback and the sources of such indiscriminately, but rather may do so in a target-dependent and motivationally-consistent manner.

Our findings also add nuance to our understanding of both primary and social reward processing. Consistent with our findings, the value people assign to primary rewards like food can be influenced by a variety of contextual and motivational factors. Although humans and animals value eating a tasty food and doing so elicits reward-related neural activity, its value diminishes as someone fills up on a food or habituates to its taste (e.g., Balleine & Dickinson, 1998; Gottfried, O’Doherty, & Dolan, 2003; Kringelbach, O’Doherty, Rolls, Andrews, 2003). Individual motives (e.g., pursuing health goals) can reduce the value of tasty foods (e.g., Hare, Camerer, & Rangel, 2009; Nook & Zaki, 2015), and the opinion of experts or the price of a wine can likewise increase or decrease the reward value of consuming such goods (e.g., Campbell-Meiklejohn, Bach, Roepstorff, Dolan, & Frith, 2010; Klucharev, Smidts, & Fernández, 2008; Plassmann, O’Doherty, Shiv, & Rangel, 2008).

A variety of social and motivational contexts likewise influence the value of social rewards. Consistent with our findings, research shows that the specific identify of social targets may influence social reward processes. For instance, people generally wish to affiliate with and enhance close others (e.g., friends, romantic partners, ingroup members) more than distant others (e.g., acquaintances, strangers, outgroup members; Hughes & Beer, 2012; Hughes, Zaki, Ambady, 2017; Ratner, Dotsch, Wigboldus, van Knippenberg, & Amodio, 2014). This amplifies the value of social experiences with close others compared to distant others. Indeed, people experience more pleasure and exhibit greater reward-related activity in response to social connection, cooperation, conformity, and vicarious reward for close versus distant others (Braams et al., 2014; Cikara, Botvinick, Fiske, 2011; Fareri, Chang, Delgado, 2015; Hackel, Zaki, Van Bavel, 2017; Hughes, Ambady, Zaki, 2017; Izuma & Adolphs, 2013; Mobbs et al., 2009; Stallen et al., 2013; Varnum et al., 2014). In fact, the value people assign to such experiences can diminish or disappear when individuals are required to interact with disliked targets or members of other groups (Cikara et al., 2011; Hein, Silani, Preuschoff, Batson, & Singer, 2010; Hughes, Ambady, Zaki, 2017; Izuma & Adolphs, 2013; Stallen et al., 2013). Individual motives and contexts can modulate the value of social rewards, just as they enhance or diminish the value and subsequent drive to attain primary rewards such as food. Thus, contextual shifts in value may be linked with changes in the likelihood of achieving desired outcomes: shifts in value can increase the likelihood of social connection with close others, and diminish the likelihood of social connection with distant or disliked others.

Together, these findings deepen our understanding of the psychological structure of social reward. Although people experience basic social signals as subjectively valuable, these social experiences do not exist in a vacuum (Hughes & Zaki, 2015; Michalska, Gardiner, & Hughes, 2018; Redcay & Warnell, 2017). Instead, a multitude of contextual and motivational features can change the extent to which pursuing a social reward will bring about a desired outcome or adaptive goal. These factors can re-shape the ultimate functionality of social signals and experiences, which can then amplify or diminish their value. Our findings suggest that the value of social signals—such as social approval—is defined by the extent to which they bring us closer to people we care about (e.g., close others). Future research that aims for a more complete picture of human sociality can thus benefit from incorporating the social contexts and motivations in which social rewards are embedded. Recontextualizing social rewards can shed new light on why people ultimately value them.

